# *Pseudomonas apudapuas* sp. nov., *Pseudomonas fontis* sp. nov., *Pseudomonas idahonensis* sp. nov., and *Pseudomonas rubra* sp. nov. isolated from in, and around, a rainbow trout farm

**DOI:** 10.1101/2022.10.04.510816

**Authors:** Todd Testerman, Jackie Varga, Hailey Donohue, Caroline Vieira Da Silva, Joerg Graf

**Affiliations:** University of Connecticut, Department of Molecular and Cell Biology, Storrs, Connecticut, USA

**Author notes:** **Corresponding author** Joerg Graf. **Repositories:** The GenBank/EMBL/DDBJ accession number for the genomes are: JAMDGP000000000 (ID291^T^); JAMDGR000000000 (ID357 ^T^); JAMDGS000000000 (ID386 ^T^); and JAMDGX000000000 (ID656 ^T^). The accession numbers for the full-length 16S rRNA gene sequences are ON555796 (ID291 ^T^); ON555782 (ID357 ^T^); ON555782 (ID386 ^T^); and ON555788 (ID656 ^T^).

**Keywords:** *Pseudomonas*, aquaculture, rainbow trout, biofilm, freshwater

## Abstract

During a large-scale bacterial culturing effort of biofilms in the vicinity of a rainbow trout aquaculture facility in Idaho, USA, ten isolates were identified as having pathogen inhibiting activity and were characterized further. These isolates were shown to be Gram negative, rod-shaped bacteria belonging to the genus *Pseudomonas*. Whole genome comparisons and multi-locus sequence analysis using four housekeeping genes (16S rDNA, *gyrA, rpoB, rpoD*) showed that these 10 isolates clustered into four distinct species groups. These comparisons also indicated that these isolates were below the established species cutoffs for the genus *Pseudomonas*. Further phenotypic characterization using API 20NE, API ZYM, and BioLog GENIII assays and chemotaxonomic analysis of cellular fatty acids were carried out. Based on the genomic, physiological, and chemotaxonomic properties of these isolates, we concluded that these strains composed four novel species of *Pseudomonas*. The proposed names are: *Pseudomonas apudapuas* sp. nov. consisting of strains ID233, ID386^T^, and ID387 with ID386^T^ (DSM 114641) as the type strain; *Pseudomonas rubra* sp. nov. consisting of strains ID291^T^, ID609, and ID1025 with ID291^T^ (DSM 114640) as the type strain; *Pseudomonas idahonensis* sp. nov. consisting of strains ID357^T^ and ID1048 with ID357^T^ (DSM 114609) as the type strain; and *Pseudomonas fontis* sp. nov. consisting of strains ID656^T^ and ID681 with ID656^T^ (DSM 114610) as the type strain.

---

The aquatic environment is a rich biome for *Pseudomonas* species and includes natural habitats as well as the built environment. Pseudomonads have incredibly diverse genomes with a suite of genetic components that allow them to survive and reproduce in a wide range of settings (1). This genus of bacteria is highly speciose, and its members have relevance as human (2–4), animal (5,6), and plant pathogens (7,8) and symbionts (9–11) as well as producers of many antimicrobial compounds that are of scientific interest (12,13).

We performed high-throughput culturing and screening of bacterial isolates found within biofilms of a rainbow trout (*Oncorhynchus mykiss*) hatchery and the upstream, natural spring environment that provides the water for this hatchery. This screening included *in vitro* testing for inhibitory activity against multiple fish and human pathogens. The inhibitory isolates were then identified using 16S rRNA gene sequencing and many were shown to belong to the genus *Pseudomonas*. With the potential for use as novel probiotics or sources of novel antimicrobial compounds, we decided to further characterize these *Pseudomonas* isolates.

## Isolation and Ecology

Isolates were obtained by swabbing biofilms on various surfaces within a rainbow trout hatchery and from swabbing rocks at the location of the source water for this facility in Idaho, USA in October of 2019. Strains ID233, ID291^T^, ID357^T^, ID386^T^, ID387, ID609, ID656^T^, and ID1048 were obtained from surfaces in rainbow trout hatchery raceways. Strains ID681 and ID1025 were obtained from rocks from Box Canyon Spring, Idaho. Samples were collected by swabbing a ∼1m long strip of the surface of interest with a sterile swab and then plated onto Reasoner’s 2A agar (R2A) (DSMZ #830), tryptic soy agar (TSA) (BD, Franklin Lakes, NJ, USA), 1:100 diluted TSA (BD), nutrient agar (BD), or Aeromonas agar (Millipore-Sigma, Burlington, MA, USA) and incubated at either 15°C or 25°C for 24-48 h (Supplementary Table 1). Individual colonies from these primary plates were then subcultured until a pure culture was obtained. All isolates were able to grow at 25°C and this temperature was used as the standard temperature going forward. Isolates were stored in their primary isolation medium with a final concentration of 20% glycerol and frozen at −80°C.

**Table 1:**
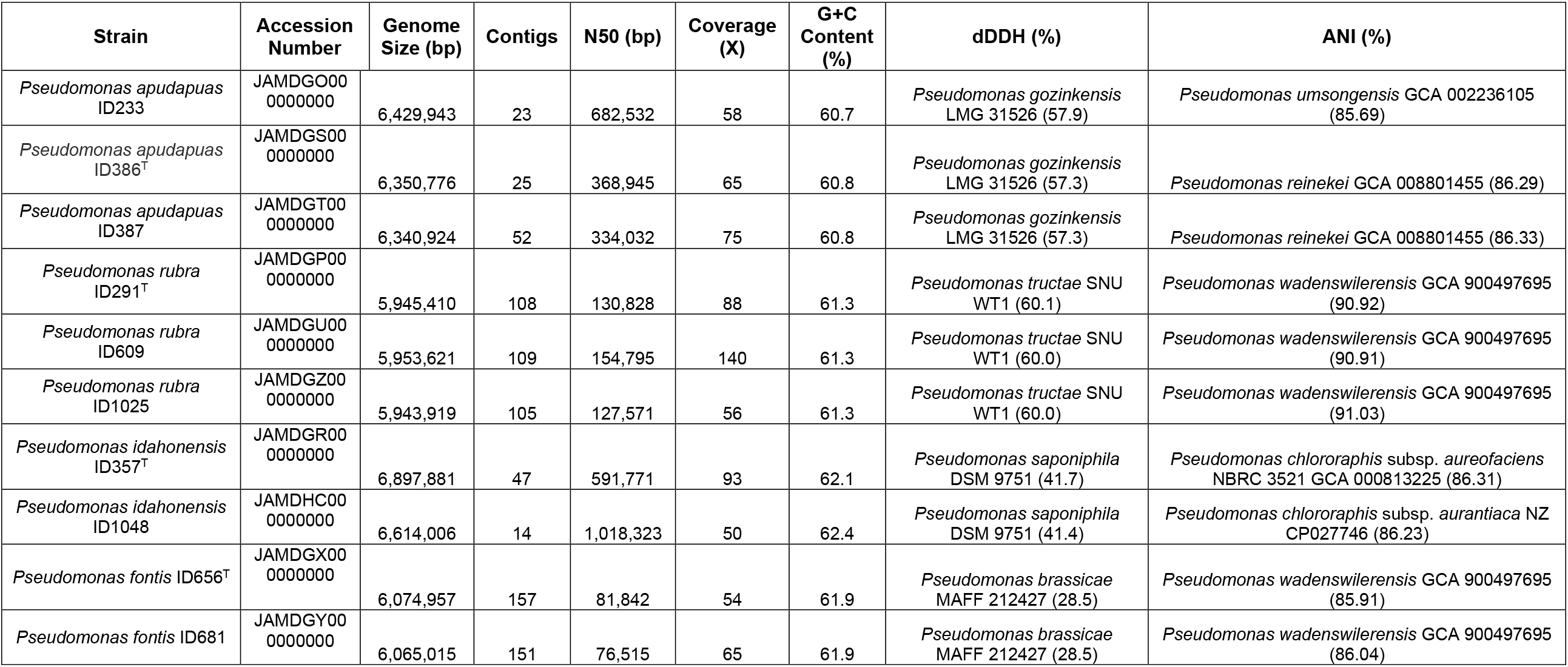
Genome Assembly Statistics and Closest Genomic Relatives. dDDH values were obtained from the Type Strain Genome

## Phylogeny

The 10 isolates were first genome sequenced, relevant genes extracted, and then compared with one another and 96 other representative *Pseudomonas* spp. First, the 16S rRNA gene sequences were used to generate a phylogeny. *Cellvibrio japonicus* was used as an outgroup and a 1,000-replicate consensus neighbor-joining tree using the Tamura-Nei genetic distance model was created using 1247 bp of the 16S rRNA gene (Supplementary Figure 1). ID357 and ID1048 clustered together on this tree but separately from other reference *Pseudomonas* spp. ID291, ID609, and ID1025 clustered together with their closest neighbor being *P. cichorii*. ID656 and ID681 clustered together with their closest neighbor being *P. putida*. ID233, ID386, and ID387 clustered together within the *P. koreensis* subgroup.

The 16S rRNA gene, while useful for broad characterization of diverse bacterial communities, struggles to attain reliable species level identification and this is especially true for genera like *Pseudomonas* (14). A more reliable approach is to concatenate multiple housekeeping genes and using a previously described strategy, this analysis was performed with these isolates (15,16). Briefly, the 16S rDNA, gyrase subunit B (*gyrB*), RNA polymerase sigma factor 70 (*rpoD*), and RNA polymerase subunit β (*rpoB*) genes from these isolates and 96 reference species were extracted from each genome, aligned, trimmed, and concatenated in this order: 16S rDNA (1,226 bp), *gyrB* (804 bp), *rpoD* (738 bp), *rpoB* (637 bp). The total length of the concatenated genes used for analysis was 3,563 bp. The same tree building parameters were used as described earlier. This tree provided far better discernment and allowed placement of these isolates into species groups (Figure 1). ID357 and ID1048 were placed within the *P. chlororaphis* subgroup of the *P. fluorescens* group. ID233, ID386, and ID387 were placed within the *P. koreensis* subgroup of the *P. fluorescens* group. ID291, ID609, and ID1025 formed a clade with ID656 and ID681 next to the *P. putida* group.

**Figure 1:**
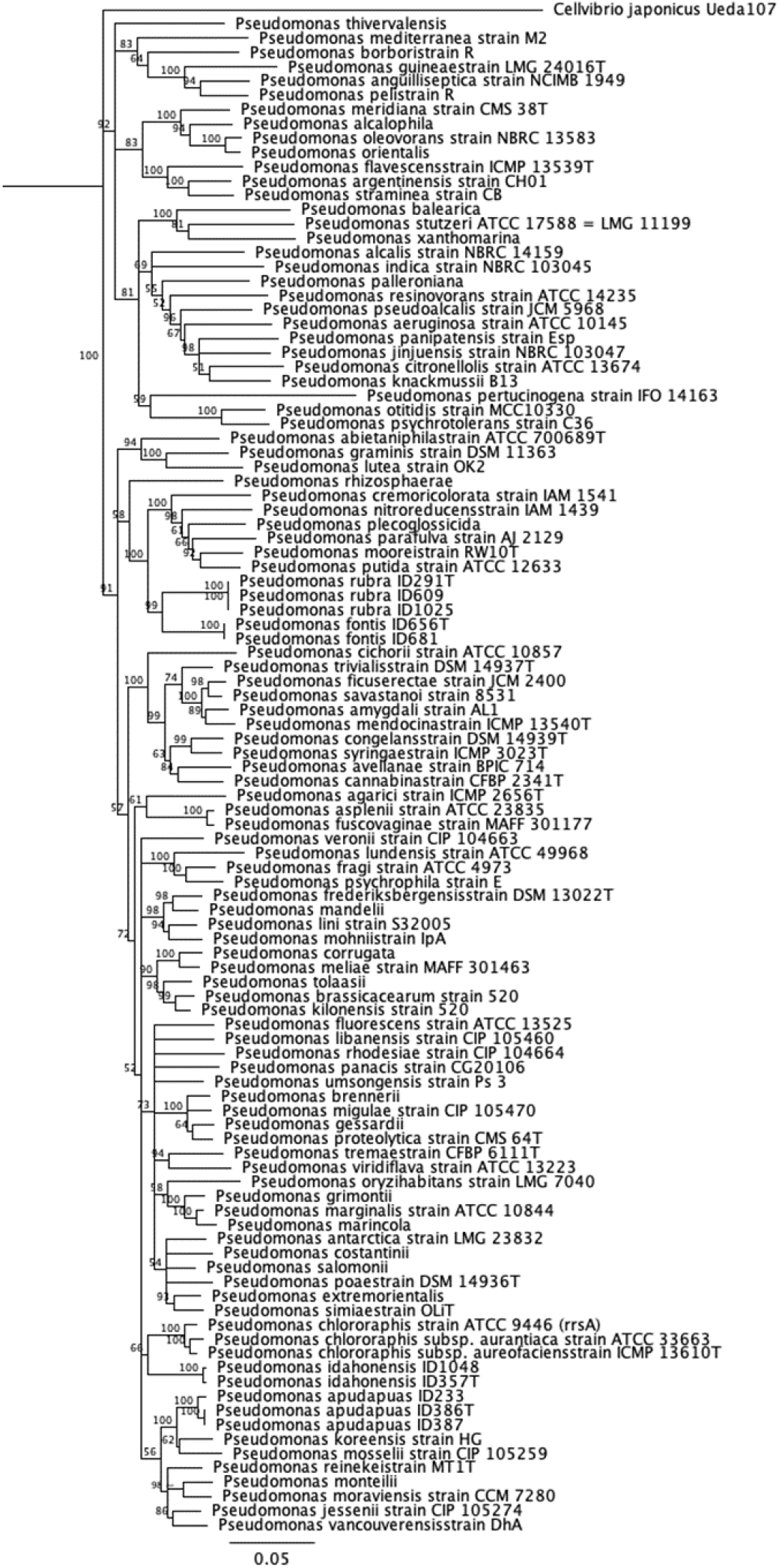
Tamura-Nei Four Gene MLSA Gene Phylogeny. 1000 replicates were used, and a consensus tree was built with branches requiring at least 50% bootstrap support.

These associations were further explored using a 100 gene Multi-Locus Sequence Typing (MLST) approach performed by autoMLST (automated Multi-Locus Species Tree) (17). Briefly, single copy housekeeping genes are automatically identified within query and reference genomes. These genes are extracted and concatenated before a phylogeny is constructed. The 30 closest reference genomes were used as determined by the program with an additional 10 genomes being manually added to represent the major *Pseudomonas* species groups. The 100 selected genes were automatically determined by autoMLST (Supplementary Table 2). The same associations as noted in the four gene MLST were observed here, albeit with better defined closest neighbors (Supplementary Figure 2). A clear distinction between our tested isolates and all reference sequences was again noted and bootstrap support were at 100/100 for almost all branches.

**Table 2:** Distinguishing Phenotypic Tests between Novel Species and Closest Genomic Relatives.

|  | **1** | **2** | **3** | **4** | **5** | **6** | **7** | **8** | **9** | **10** |
| --- | --- | --- | --- | --- | --- | --- | --- | --- | --- | --- |
| Fluorescent Pigment | - | + | - | - | + | - | ND | + | + | + |
| Growth Temperature Range | 4-37 | 4-39 | 4-37 | 4-33 | 5-32 | 4-39 | 4-45 | 4-35 | 4-37 | 4-36 |
| **API ZYM** |  |  |  |  |  |  |  |  |  |  |
| Alkaline Phosphatase | - | + | + | - | - | + | ND | ND | - | ND |
| **API 20NE** |  |  |  |  |  |  |  |  |  |  |
| Reduction of Nitrates | - | - | + | - | - | - | + | - | + | - |
| Fermentation of Glucose | - | - | - | - | - | - | + | - | - | - |
| Assimilation of N-Acetyl-Glucosamine | + | + | + | - | + | - | + | - | + | - |
| Assimilation of Adipic Acid | - | + | - | - | - | - | ND | - | - | - |
| Assimilation of Trisodium Citrate | + | + | - | + | + | + | ND | + | + | + |
| **Assimilation of (Biolog GEN III):** |  |  |  |  |  |  |  |  |  |  |
| Dextrin | - | - | - | - | - | + | ND | - | ND | + |
| N-Acetyl-D-Glucosamine | - | - | - | - | + | + | + | - | + | + |
| D-Saccharic Acid | + | - | - | - | + | - | - | - | - | - |
| Inosine | w | w | - | w | - | + | + | + | + | + |
| p-Hydroxy-Phenylacetic Acid | - | + | + | - | - | + | + | + | + | - |
| D-Malic Acid | - | - | - | + | - | - | ND | + | ND | - |
| D-Serine | - | - | - | + | + | + | + | - | + | - |
| Sucrose | - | + | - | - | - | + | - | - | + | - |
| Mucic Acid | + | - | - | + | + | ND | ND | - | ND | ND |
| Methyl Pyruvate | - | - | - | + | + | + | ND | + | + | ND |
| α-Keto-Butyric Acid | - | - | - | w | - | + | ND | + | - | ND |
| L-Alanine | w | - | - | + | + | + | ND | + | + | + |
| L-Serine | + | - | w | + | + | - | ND | + | + | + |
| Glycerol | - | w | - | + | + | + | ND | + | + | + |
| D-Fructose-6-Phosphate | - | + | - | - | + | - | ND | ND | - | ND |
| Glucuronamide | + | w | - | w | - | - | + | ND | - | ND |
| L-Aspartic Acid | - | w | - | + | + | + | ND | + | + | ND |
| D-Galacturonic Acid | - | - | - | - | - | - | + | + | - | - |
| D-Glucuronic Acid | - | - | - | - | - | - | + | ND | - | + |
| D-Fructose | - | w | - | - | - | + | ND | + | + | + |
| L-Galactonic Acid-γ-Lactone | - | w | - | - | - | - | ND | + | - | ND |
| Quinic Acid | + | + | - | - | + | + | ND | + | + | + |
| α-Hydroxy-Butyric Acid | - | - | - | - | - | - | ND | + | - | - |
| Tween 40 | + | + | w | + | w | + | ND | - | + | ND |

1- *Pseudomonas apudapuas* ID386<sup>T</sup>, 2- *Pseudomonas idahonensis* ID357<sup>T</sup>, 3- *Pseudomonas rubra* ID291<sup>T</sup>, 4- *Pseudomonas fontis* ID656<sup>T</sup>, 5- *Pseudomonas* *gozinkensis* LMG 31526, 6- *Pseudomonas saponiphila* DSM 9751, 7- *Pseudomonas tructae* KCTC 72265, 8- *Pseudomonas brassicae* MAFF 212427, 9-
*Pseudomonas donghuensis* DSM 101685, 10- *Pseudomonas protegens* DSM 19095. Symbols: -, negative reaction; w, weak reaction; +, positive reaction; ND, no data; ANI, Average Nucleotide Identity; dDDH, digital DNA-DNA Hybridization.

## Genome Features

The genomes of the 10 isolates were prepared using the Illumina DNA Prep kit (Illumina, San Diego, CA, USA) and sequenced using 250×250 bp paired-end sequencing on an Illumina MiSeq (Illumina). The genomes were processed and assembled using CLC (version 22.0, Qiagen, Hilden, Germany) before being annotated by the Prokaryotic Genome Annotation Pipeline (version 6.0) (18) and deposited on NCBI. The assembled genomes were then analyzed, and digital DNA-DNA hybridization (dDDH) and Average Nucleotide Identity (ANI) percentages were calculated. Table 1 displays genome statistics and the highest percentage reference species identities for each novel strain.

These results supported the earlier conclusions based on MLST and also established that these strains are novel genomic species within the genus *Pseudomonas*. All sequenced genomes were below the 95% ANI similarity threshold and the 70% dDDH similarity threshold. A genome-based phylogeny can be seen in Supplementary Figure 3.

## Physiology and Chemotaxonomy

For morphologic characterization, the four type strains were grown in R2A broth at 25°C for 24h and visualized using transmission electron microscopy (TEM) (Supplementary Figure 4). All strains were rod-shaped with one or more polar flagella and were 1-3 μm in length and 0.8 μm in diameter. On R2A, colonies were generally 1-2mm in diameter and white/off-white with the exception of ID291^T^, ID609, and ID1025, which were pink/red in color. All strains stained Gram-negative and were both catalase and oxidase positive.

Physiological examination included temperature, pH, and salinity range testing using Luria-Bertani (LB) medium incubated at 25°C for 24-48h. Pyocyanin and fluorescent pigment production were also determined using King A (Millipore-Sigma) and King B (19) media, respectively, incubated at 25°C for 5 days. API ZYM (bioMérieux, Marcy-l’Étoile, France) and API 20NE (bioMérieux) strips were used to examine enzymatic and metabolic potentials of all strains. Biolog GEN III plates (Biolog, Hayward, CA, USA) were also used (only for strains ID233, ID291^T^, ID357^T^, ID386^T^, ID609, ID656^T^, and ID1048) to further characterize metabolic potential and chemical sensitivity (Supplementary Table 3). Additionally, closely related reference strains, as determined from phylogenetic and genomic comparisons, were included for many of these tests. A summary of the traits that differentiate these proposed novel strains from one another as well as other closely related *Pseudomonas* spp. are presented in Table 2.

Cellular fatty acid composition profiling of the four proposed type strains was carried out by DSMZ Services (Leibniz-Institut DSMZ, Braunschweig, Germany) using fatty acid methyl ester (FAME) gas chromatography with peak identities confirmed by gas chromatography-mass spectrometry (GC-MS). Strains were grown on R2A medium and incubated for 24-48h before profiling. The main fatty acids were C_16:1_ ω7c (20.4-39.6%), C_16:0_ (22.7-28.6%), and C_18:1_ ω7c (11.2-19.3%) (Supplementary Table 4). All strains contained the characteristic *Pseudomonas* fatty acids C_10:0 3-OH_ (5.1-11%), C_12:0_ (1.5-2.5%), and C_12:0 2-OH_ (4.7-6.5%) (20).

## Description of *Pseudomonas apudapuas* sp. nov

*Pseudomonas apudapuas* (a.pud.a.pu’as, L. pref. *apud* – among; L. fem. n. acc. *apuas* - young fish; N.L fem. adj. *apudapuas* - among young fish, referring to its isolation from a fish hatchery).

Cells are Gram-negative, rod-shaped, aerobic, oxidase-positive, catalase-positive, possess lophotrichous flagella, and measure approximately 2 μm in length and 0.8 μm in diameter. Colonies are 1-2 mm in size, white, and well-defined on R2A. This species grows between 4-37°C (with an optimum being 25°C) but does not grow at 38°C. This species grows within a NaCl range of 0-5% and a pH range between 6-8. Pyocyanin is not produced on King A agar and fluorescent pigment is not produced on King B agar. API 20NE results show this species to be positive for arginine dihydrolase and gelatin hydrolysis. This strain is able to assimilate D-glucose, L-arabinose, D-mannose, D-mannitol, N-acetyl-glucosamine, potassium gluconate, capric acid, malic acid, and trisodium citrate. All other tests on the API 20NE strip were deemed to have a negative result. API ZYM results show this species to have enzymatic activity for esterase (C4), esterase lipase (C8), leucine arylamidase, acid phosphatase, and naphthol-AS-BI-phosphohydrolase. All other tests on the API ZYM strip were deemed to have a negative result. Biolog GENIII plates results showed this species to be able to utilize D-glucose, D-mannose, D-galactose, D-fucose, L-fucose, inosine, L-alanine, L-arginine, L-pyroglutamic acid, L-serine, L-galactonic acid-γ-lactone, D-gluconic acid, glucuronamide, mucic acid, quinic acid, D-saccharic acid, L-lactic acid, citric acid, α-keto-glutaric acid, L-malic acid, bromo-succinic acid, Tween 40, γ-amino-n-butyric acid, β-hydroxy-butyric acid, propionic acid, and acetic acid. This species is able to grow in the presence of 1% sodium lactate, fusidic acid, troleandomycin, rifamycin SV, lincomycin, guanidine hydrochloride, Niaproof, vancomycin, tetrazolium violet, tetrazolium blue, nalidixic acid, lithium chloride, potassium tellurite, aztreonam, and sodium bromate. All other tests on the Biolog GENIII plate were deemed to have a negative result. The predominant cellular fatty acids are C_16:1_ ω7c (39.6%), C_16:0_ (28.6%), C_18:1_ ω7c (11.2%), C_12:0 2-OH_ (6.5%), C_10:0 3-OH_ (5.1%), C_12:0 3-OH_ (5.1%), C_12:0_ (1.5%), and C_17:0_ cyclo ω7c (1.2%). This species can be distinguished from its closest related species (represented by LMG 31526) by its ability to grow at 37°C, a relatively low percentage of C_17:0_ cyclo fatty acid (1.2% vs. 17.7% [LMG 31526]), utilization of glucuronamide, and its inability to utilize D-serine, methyl pyruvate, glycerol, D-fructose-6-phosphate, and L-aspartic acid.

The type strain is ID386^T^ (DSM 114641) and was isolated from a biofilm formed on the surface of a wall within a rainbow trout hatchery building. The genomic DNA G+C content of the type strain is 60.8 mol%. The draft genome for strain ID386^T^ has been deposited in the DDBJ/ENA/GenBank databases under the accession number JAMDGS000000000. The version described in this paper is the first version. The 16S rRNA gene sequence for strain ID386^T^ is available under the accession number ON555786.

## Description of *Pseudomonas idahonensis* sp. nov

*Pseudomonas idahonensis* (i.da.ho.nen’sis, N.L. neut. n. *idaho* – Idaho, USA; L. suff. *nensis* – from; N.L. neut. adj. *idahonensis* – from Idaho, referring to the U.S. state where this species was isolated).

Cells are Gram-negative, rod-shaped, aerobic, oxidase-positive, catalase-positive, possess lophotrichous flagella, and measure approximately 1.5 μm in length and 0.8 μm in diameter. Colonies are 1-2 mm in size, opaque white, and well-defined on R2A. This species grows between 4-39°C (with an optimum being 25°C) but does not grow at 40°C. This species grows within a NaCl range of 0-5% and a pH range between 6-8. Pyocyanin is not produced on King A agar, but fluorescent pigment is produced on King B agar. API 20NE results show this species to be positive for arginine dihydrolase and gelatin hydrolysis. This strain is able to assimilate D-glucose, D-mannose, D-mannitol, N-acetyl-glucosamine, potassium gluconate, capric acid, adipic acid, malic acid, trisodium citrate, and phenylacetic acid. All other tests on the API 20NE strip were deemed to have a negative result. API ZYM results show this species to have enzymatic activity for alkaline phosphatase, esterase (C4), esterase lipase (C8), leucine arylamidase, acid phosphatase, and naphthol-AS-BI-phosphohydrolase. All other tests on the API ZYM strip were deemed to have a negative result. Biolog GENIII plates results showed this species to be able to utilize D-trehalose, sucrose, D-glucose, D-mannose, D-fructose, D-fucose, L-fucose, inosine, myo-inositol, glycerol, D-fructose-6-phosphate, L-arginine, L-glutamic acid, L-histidine, pectin, D-gluconic acid, glucuronamide, quinic acid, p-hydroxy-phenylacetic acid, L-lactic acid, citric acid, α-keto-glutaric acid, L-malic acid, bromo-succinic acid, Tween 40, γ-amino-n-butyric acid, β-hydroxy-butyric acid, propionic acid, and acetic acid. This species is able to grow in the presence of 1% sodium lactate, fusidic acid, D-serine, troleandomycin, rifamycin SV, minocycline, lincomycin, guanidine hydrochloride, Niaproof, vancomycin, tetrazolium violet, tetrazolium blue, nalidixic acid, potassium tellurite, aztreonam, and sodium bromate. All other tests on the Biolog GENIII plate were deemed to have a negative result. The predominant cellular fatty acids are C_16:0_ (22.8%), C_16:1_ ω7c (21.7%), C_18:1_ ω7c (19.3%), C_10:0 3-OH_ (7.5%), C_12:0 3-OH_ (7.2%), C_12:0 2-OH_ (5.1%), C_17:0_ cyclo ω7c (4.6%), C_16:1_ ω7t (3.3%), C_12:1 3-OH_ ω7c (3.0%), C_12:0_ (2.0%), and C_14:0 3-OH_ (1.2%). This species can be distinguished from its closest related species (represented by DSM 9751) by its production of fluorescent pigment on King B agar, its assimilation of adipic acid, and its inability to utilize dextrin, D-serine, methyl pyruvate, α-keto-butyric acid, and L-alanine.

The type strain is ID357^T^ (DSM 114609) and was isolated from a biofilm formed on the surface of a wall within a rainbow trout hatchery building. The genomic DNA G+C content of the type strain is 62.1 mol%. The draft genome for strain ID357^T^ has been deposited in the DDBJ/ENA/GenBank databases under the accession number JAMDGR000000000. The version described in this paper is the first version. The 16S rRNA gene sequence for strain ID357^T^ is available under the accession number ON555782.

## Description of *Pseudomonas rubra* sp. nov

*Pseudomonas rubra* (ru’bra, L. fem. adj. *rubra* – red, referring to the colony color).

Cells are Gram-negative, rod-shaped, aerobic, oxidase-positive, catalase-positive, possess lophotrichous flagella, and measure approximately 2 μm in length and 0.8 μm in diameter. Colonies are 1-2 mm in size, opaque white with a red center, and are not well-defined on R2A. This species grows between 4-37°C (with an optimum being 25°C) but does not grow at 38°C. This species grows within a NaCl range of 0-5% and a pH range between 6-9. Pyocyanin is not produced on King A agar and fluorescent pigment is not produced on King B agar. API 20NE results show this species to be positive for nitrate reduction, arginine dihydrolase and gelatin hydrolysis. This strain is able to assimilate D-glucose, N-acetyl-glucosamine, potassium gluconate, capric acid, and malic acid,. All other tests on the API 20NE strip were deemed to have a negative result. API ZYM results show this species to have enzymatic activity for alkaline phosphatase, esterase (C4), leucine arylamidase, valine arylamidase, cystine arylamidase, acid phosphatase, and naphthol-AS-BI-phosphohydrolase. All other tests on the API ZYM strip were deemed to have a negative result. Biolog GENIII plates results showed this species to be able to utilize D-glucose, L-glutamic acid, L-histidine, L-serine, D-gluconic acid, p-hydroxy-phenylacetic acid, L-lactic acid, L-malic acid, Tween 40, β-hydroxy-butyric acid, and acetic acid. This species is able to grow in the presence of 1% sodium lactate, fusidic acid, D-serine, troleandomycin, rifamycin SV, minocycline, lincomycin, guanidine hydrochloride, Niaproof, vancomycin, tetrazolium violet, tetrazolium blue, nalidixic acid, potassium tellurite, and aztreonam. All other tests on the Biolog GENIII plate were deemed to have a negative result. The predominant cellular fatty acids are C_16:0_ (23.9%), C_16:1_ ω7c (22.2%), C_18:1_ ω7c (17.2%), C_10:0 3-OH_ (11.0%), C_16:1_ ω7t (6.6%), C_12:0 3-OH_ (6.1%), C_12:0 2-OH_ (5.4%), C_12:1 3-OH_ ω7c (2.4%), C_17:0_ cyclo ω7c (1.9%), and C_12:0_ (1.6%). This species can be phenotypically distinguished from its closest related species (represented by KCTC 72265) by its inability to grow above 39°C, inability to grow at NaCl concentrations above 5%, its inability to ferment glucose, and its inability to utilize inosine, D-serine, glucuronamide, D-galacturonic acid, and D-glucuronic acid.

The type strain is ID291^T^ (DSM 114640) and was isolated from a biofilm formed on the surface of a wall within a rainbow trout hatchery building. The genomic DNA G+C content of the type strain is 61.3 mol%. The draft genome for strain ID291^T^ has been deposited in the DDBJ/ENA/GenBank databases under the accession number JAMDGP000000000. The version described in this paper is the first version. The 16S rRNA gene sequence for strain ID291^T^ is available under the accession number ON555796.

## Description of *Pseudomonas fontis* sp. nov

*Pseudomonas fontis* (fon’tis, L. gen. n. *fontis* – of a spring, referring to this species’ isolation from a freshwater spring).

Cells are Gram-negative, rod-shaped, aerobic, oxidase-positive, catalase-positive, possess lophotrichous flagella, and measure approximately 2 μm in length and 0.8 μm in diameter. Colonies are 1-2 mm in size, white, and well-defined on R2A. This species grows between 4-33°C (with an optimum being 25°C) but does not grow at 34°C. This species grows within a NaCl range of 0-5% and a pH range between 6-8. Pyocyanin is not produced on King A agar and fluorescent pigment is not produced on King B agar. API 20NE results show this species to be positive for arginine dihydrolase and gelatin hydrolysis. This strain is able to assimilate D-glucose, potassium gluconate, capric acid, malic acid, and trisodium citrate. All other tests on the API 20NE strip were deemed to have a negative result. API ZYM results show this species to have enzymatic activity for esterase (C4), leucine arylamidase, and naphthol-AS-BI-phosphohydrolase. All other tests on the API ZYM strip were deemed to have a negative result. Biolog GENIII plates results showed this species to be able to utilize D-glucose, D-fucose, L-fucose, inosine, glycerol, D-serine, L-alanine, L-arginine, L-aspartic acid, L-glutamic acid, L-histidine, L-serine, D-gluconic acid, glucuronamide, mucic acid, methyl pyruvate, L-lactic acid, citric acid, α-keto-glutaric acid, D-malic acid, L-malic acid, bromo-succinic acid, Tween 40, γ-amino-n-butyric acid, β-hydroxy-butyric acid, α-keto-butyric acid, propionic acid, and acetic acid. This species is able to grow in the presence of 1% sodium lactate, fusidic acid, D-serine, troleandomycin, rifamycin SV, lincomycin, guanidine hydrochloride, Niaproof, vancomycin, tetrazolium violet, tetrazolium blue, nalidixic acid, potassium tellurite, aztreonam, and sodium bromate. All other tests on the Biolog GENIII plate were deemed to have a negative result. The predominant cellular fatty acids are C_16:0_ (22.7%), C_16:1_ ω7c (20.4%), C_18:1_ ω7c (19.3%), C_10:0 3-OH_ (9.2%), C_17:0_ cyclo ω7c (6.4%), C_12:0 3-OH_ (6.0%), C_12:0 2-OH_ (4.7%), C_16:1_ ω7t (4.2%), C_12:1 3-OH_ ω7c (3.1%), and C_12:0_ (2.5%). This species can be phenotypically distinguished from its closest related species (represented by MAFF 212427) by its inability to grow at 35°C, its utilization of D-serine, mucic acid, and Tween 40, and its inability to utilize p-hydroxy-phenylacetic acid, D-galacturonic acid, D-fructose, L-galactonic acid-γ-lactone, quinic acid, and α-hydroxy-butyric acid.

The type strain is ID656^T^ (DSM 114610) and was isolated from a biofilm formed on the surface of a wall within a rainbow trout hatchery building. The genomic DNA G+C content of the type strain is 60.8 mol%. The draft genome for strain ID656^T^ has been deposited in the DDBJ/ENA/GenBank databases under the accession number JAMDGX000000000. The version described in this paper is the first version. The 16S rRNA gene sequence for strain ID656^T^ is available under the accession number ON555788.

## AUTHOR STATEMENTS

### 1.6 Authors and contributors

Conceptualization – JG, TT

Data curation – TT, JV

Formal Analysis – TT, JV

Funding acquisition - JG

Investigation – TT, JG, JV, CVDS

Methodology - TT

Project administration – JG

Resources - JG

Software – JG

Supervision – JG, TT

Validation – TT, JV

Visualization – TT, CVDS

Writing – original draft - TT

Writing – review & editing – TT, JG, CVDS

## 1.9 Acknowledgements

We thank the UConn Microbial Analysis, Resources, and Services (MARS) facility for sequencing the genomes of these strains. We thank the Biosciences Electron Microscopy Facility of the University of Connecticut and specifically Dr. Xuanhao Sun for assistance in obtaining TEM images.

## ABBREVIATIONS

*<u>Instructions – please delete before submission</u>*. *Please include any non-standard abbreviations*.

HKY: Hasegawa, Kishino and Yano
MLST: Multi-Locus Sequence Typing
dDDH: digital DNA-DNA Hybridization
ANI: Average Nucleotide Identity
TEM: Transmission Electron Microscopy

## REFERENCES

1. Silby MW, Winstanley C, Godfrey SAC, Levy SB, Jackson RW. Pseudomonas genomes: diverse and adaptable. FEMS Microbiology Reviews [Internet]. 2011 Jul 1 [cited 2022 Jun 23];35(4):652–80. Available from: https://academic.oup.com/femsre/article/35/4/652/630861

2. Moradali MF, Ghods S, Rehm BHA. Pseudomonas aeruginosa lifestyle: A paradigm for adaptation, survival, and persistence. Frontiers in Cellular and Infection Microbiology. 2017 Feb 15;7(FEB):39.

3. Iglewski BH. Pseudomonas. 1996.

4. Kim SE, Park SH, Park HB, Park KH, Kim SH, Jung SI, et al. Nosocomial Pseudomonas putida Bacteremia: High Rates of Carbapenem Resistance and Mortality. Chonnam Medical Journal [Internet]. 2012 [cited 2022 May 27];48(2):91. Available from: /pmc/articles/PMC3434797/

5. Park H, Hong M, Hwang S, Park Y, Kwon K, Yoon J, et al. Characterisation of Pseudomonas aeruginosa related to bovine mastitis. Acta Vet Hung [Internet]. 2014 Mar 1 [cited 2022 May 26];62(1):1–12. Available from: https://pubmed.ncbi.nlm.nih.gov/24334080/

6. Bobrov AG, Getnet D, Swierczewski B, Jacobs A, Medina-Rojas M, Tyner S, et al. Evaluation of Pseudomonas aeruginosa pathogenesis and therapeutics in military-relevant animal infection models. APMIS [Internet]. 2021 [cited 2022 May 27]; Available from: https://onlinelibrary.wiley.com/doi/full/10.1111/apm.13119

7. Xin XF, Kvitko B, He SY. Pseudomonas syringae: what it takes to be a pathogen. Nature Reviews Microbiology 2018 16:5 [Internet]. 2018 Feb 26 [cited 2022 May 26];16(5):316–28. Available from: https://www.nature.com/articles/nrmicro.2018.17

8. Höfte M, de Vos P. Plant pathogenic species. Plant-Associated Bacteria [Internet]. 2007 [cited 2022 May 26];507–33. Available from: https://link.springer.com/chapter/10.1007/978-1-4020-4538-7_14

9. Gera Hol WH, Martijn Bezemer T, Biere A. Getting the ecology into interactions between plants and the plant growth-promoting bacterium Pseudomonas fluorescens. Frontiers in Plant Science. 2013 Apr 10;4(APR):81.

10. Arrebola E, Tienda S, Vida C, de Vicente A, Cazorla FM. Fitness features involved in the biocontrol interaction of pseudomonas chlororaphiswith host plants: The case study of PcPCL1606. Frontiers in Microbiology. 2019;10(APR):719.

11. Waghunde RR, Sabalpara AN. Impact of Pseudomonas spp. on Plant Growth, Lytic Enzymes and Secondary Metabolites Production. Frontiers in Agronomy. 2021 Dec 14;3:98.

12. Shahid I, Malik KA, Mehnaz S. A decade of understanding secondary metabolism in Pseudomonas spp. for sustainable agriculture and pharmaceutical applications. Environmental Sustainability 2018 1:1 [Internet]. 2018 May 24 [cited 2022 May 27];1(1):3–17. Available from: https://link.springer.com/article/10.1007/s42398-018-0006-2

13. Gross H, Loper JE. Genomics of secondary metabolite production by Pseudomonas spp. Nat Prod Rep [Internet]. 2009 [cited 2022 May 27];26(11):1408–46. Available from: https://pubmed.ncbi.nlm.nih.gov/19844639/

14. Rajwar A, Sahgal M. Phylogenetic relationships of fluorescent pseudomonads deduced from the sequence analysis of 16S rRNA, Pseudomonas-specific and rpoD genes. 3 Biotech [Internet]. 2016 Dec 1 [cited 2022 Jun 4];6(1):1–10. Available from: https://link.springer.com/article/10.1007/s13205-016-0386-x

15. Gomila M, Peña A, Mulet M, Lalucat J, García-Valdés E. Phylogenomics and systematics in Pseudomonas. Frontiers in Microbiology. 2015;6(MAR):214.

16. Mulet M, Lalucat J, García-Valdés E. DNA sequence-based analysis of the Pseudomonas species. Environmental Microbiology [Internet]. 2010 Jun 1 [cited 2022 Jun 4];12(6):1513–30. Available from: https://onlinelibrary.wiley.com/doi/full/10.1111/j.1462-2920.2010.02181.x

17. Alanjary M, Steinke K, Ziemert N. AutoMLST: an automated web server for generating multi-locus species trees highlighting natural product potential. Nucleic Acids Research [Internet]. 2019 Jul 2 [cited 2022 Jun 4];47(W1):W276–82. Available from: https://academic.oup.com/nar/article/47/W1/W276/5475077

18. Tatusova T, Dicuccio M, Badretdin A, Chetvernin V, Nawrocki EP, Zaslavsky L, et al. NCBI prokaryotic genome annotation pipeline. Nucleic Acids Research [Internet]. 2016 Aug 8 [cited 2022 Jun 4];44(14):6614. Available from: /pmc/articles/PMC5001611/

19. King EO, Ward MK, Raney DE. Two simple media for the demonstration of pyocyanin and fluorescin. J Lab Clin Med [Internet]. 1954 Aug [cited 2022 Jun 6];44(2):301–7. Available from: https://pubmed.ncbi.nlm.nih.gov/13184240/

20. Palleroni N, Garrity G, Bell J, Lilburn T. Bergey’s Manual® of Systematic Bacteriology. Vol. 2, Bergey’s Manual® of Systematic Bacteriology. Springer US; 2005. 323–354 p.

21. Meier-Kolthoff JP, Göker M. TYGS is an automated high-throughput platform for state-of-the-art genome-based taxonomy. Nature Communications 2019 10:1 [Internet]. 2019 May 16 [cited 2022 Jun 5];10(1):1–10. Available from: https://www.nature.com/articles/s41467-019-10210-3

22. Rodriguez-R LM, Gunturu S, Harvey WT, Rosselló-Mora R, Tiedje JM, Cole JR, et al. The Microbial Genomes Atlas (MiGA) webserver: taxonomic and gene diversity analysis of Archaea and Bacteria at the whole genome level. Nucleic Acids Research [Internet]. 2018 Jul 7 [cited 2022 Jun 5];46(Web Server issue):W282. Available from: /pmc/articles/PMC6031002/

