## Supplementary for "*Pseudomonas apudapuas* sp. nov., *Pseudomonas fontis* sp. nov., *Pseudomonas idahonensis* sp. nov., and *Pseudomonas rubra* sp. nov. isolated from in, and around, a rainbow trout farm"

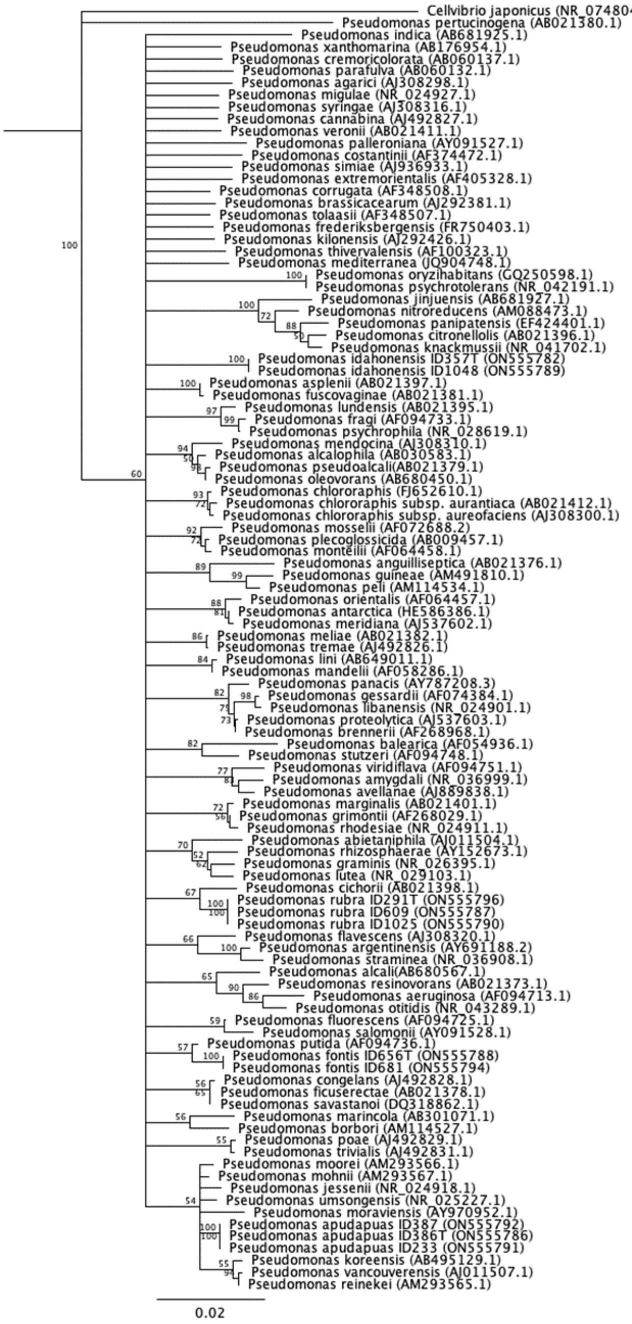
Cellvibrio japonicus (NR_074804.1)
Pseudomonas pertucinogena (AB021380.1)
Pseudomonas indica (AB681925.1)
Pseudomonas xanthomarina (AB176954.1)
Pseudomonas cremoricolorata (AB060137.1)
Pseudomonas parafulva (AB060132.1)
Pseudomonas agarici (AJ308298.1)
Pseudomonas migulae (NR_024927.1)
Pseudomonas syringae (AJ308316.1)
Pseudomonas cannabina (AJ492827.1)
Pseudomonas veronii (AB021411.1)
Pseudomonas palleroniana (AY091527.1)
Pseudomonas costantinii (AF374472.1)
Pseudomonas simiae (AJ936933.1)
Pseudomonas extremorientalis (AF405328.1)
Pseudomonas corrugata (AF348508.1)
Pseudomonas brassicacearum (AJ292381.1)
Pseudomonas tolaasii (AF348507.1)
Pseudomonas frederiksbergensis (FR750403.1)
Pseudomonas kilonensis (AJ292426.1)
Pseudomonas thivervalensis (AF100323.1)
Pseudomonas mediterranea (JQ904748.1)
Pseudomonas oryzihabitans (GQ250598.1)
Pseudomonas psychrotolerans (NR_042191.1)
Pseudomonas jinjuensis (AB681927.1)
Pseudomonas nitroreducens (AM088473.1)
Pseudomonas panipatensis (EF424401.1)
Pseudomonas citronellolis (AB021396.1)
Pseudomonas knackmussii (NR_041702.1)
Pseudomonas idahonensis ID357T (ON555782)
Pseudomonas idahonensis ID1048 (ON555789)
Pseudomonas asplenii (AB021397.1)
Pseudomonas fuscovaginae (AB021381.1)
Pseudomonas lundensis (AB021395.1)
Pseudomonas fragi (AF094733.1)
Pseudomonas psychrophila (NR_028619.1)
Pseudomonas mendocina (AJ308310.1)
Pseudomonas alcalophila (AB030583.1)
Pseudomonas pseudoalcali(AB021379.1)
Pseudomonas oleovorans (AB680450.1)
Pseudomonas chlororaphis (FJ652610.1)
Pseudomonas chlororaphis subsp. aurantiaca (AB021412.1)
Pseudomonas chlororaphis subsp. aureofaciens (AJ308300.1)
Pseudomonas mosselii (AF072688.2)
Pseudomonas plecoglossicida (AB009457.1)
Pseudomonas monteilii (AF064458.1)
Pseudomonas anguilliseptica (AB021376.1)
Pseudomonas guineae (AM491810.1)
Pseudomonas peli (AM114534.1)
Pseudomonas orientalis (AF064457.1)
Pseudomonas antarctica (HE586386.1)
Pseudomonas meridiana (AJ537602.1)
Pseudomonas meliae (AB021382.1)
Pseudomonas tremae (AJ492826.1)
Pseudomonas lini (AB649011.1)
Pseudomonas mandelii (AF058286.1)
Pseudomonas panacis (AY787208.3)
Pseudomonas gessardii (AF074384.1)
Pseudomonas libanensis (NR_024901.1)
Pseudomonas proteolytica (AJ537603.1)
Pseudomonas brennerii (AF268968.1)
Pseudomonas balearica (AF054936.1)
Pseudomonas stutzeri (AF094748.1)
Pseudomonas viridiflava (AF094751.1)
Pseudomonas amygdali (NR_036999.1)
Pseudomonas avellanae (AJ889838.1)
Pseudomonas marginalis (AB021401.1)
Pseudomonas grimontii (AF268029.1)
Pseudomonas rhodesiae (NR_024911.1)
Pseudomonas abietaniphila (AJ011504.1)
Pseudomonas rhizosphaerae (AY152673.1)
Pseudomonas graminis (NR_026395.1)
Pseudomonas lutea (NR_029103.1)
Pseudomonas cichorii (AB021398.1)
Pseudomonas rubra ID291T (ON555796)
Pseudomonas rubra ID609 (ON555787)
Pseudomonas rubra ID1025 (ON555790)
Pseudomonas flavescens (AJ308320.1)
Pseudomonas argentinensis (AY691188.2)
Pseudomonas straminea (NR_036908.1)
Pseudomonas alcali(AB680567.1)
Pseudomonas resinovorans (AB021373.1)
Pseudomonas aeruginosa (AF094713.1)
Pseudomonas otitidis (NR_043289.1)
Pseudomonas fluorescens (AF094725.1)
Pseudomonas salomonii (AY091528.1)
Pseudomonas putida (AF094736.1)
Pseudomonas fontis ID656T (ON555788)
Pseudomonas fontis ID681 (ON555794)
Pseudomonas congelans (AJ492828.1)
Pseudomonas ficuserectae (AB021378.1)
Pseudomonas savastanoi (DQ318862.1)
Pseudomonas marincola (AB301071.1)
Pseudomonas borbori (AM114527.1)
Pseudomonas poae (AJ492829.1)
Pseudomonas trivialis (AJ492831.1)
Pseudomonas moorei (AM293566.1)
Pseudomonas mohnii (AM293567.1)
Pseudomonas jessenii (NR_024918.1)
Pseudomonas umsongensis (NR_025227.1)
Pseudomonas moraviensis (AY970952.1)
Pseudomonas apudapuas ID387 (ON555792)
Pseudomonas apudapuas ID386T (ON555786)
Pseudomonas apudapuas ID233 (ON555791)
Pseudomonas koreensis (AB495129.1)
Pseudomonas vancouverensis (AJ011507.1)
Pseudomonas reinekei (AM293565.1)
0.02

**Supplementary Figure 1: Tamura-Nei Full-Length 16S rRNA Gene Phylogeny.** 1000 replicates were used, and a consensus tree was built with branches requiring at least 50% bootstrap support.

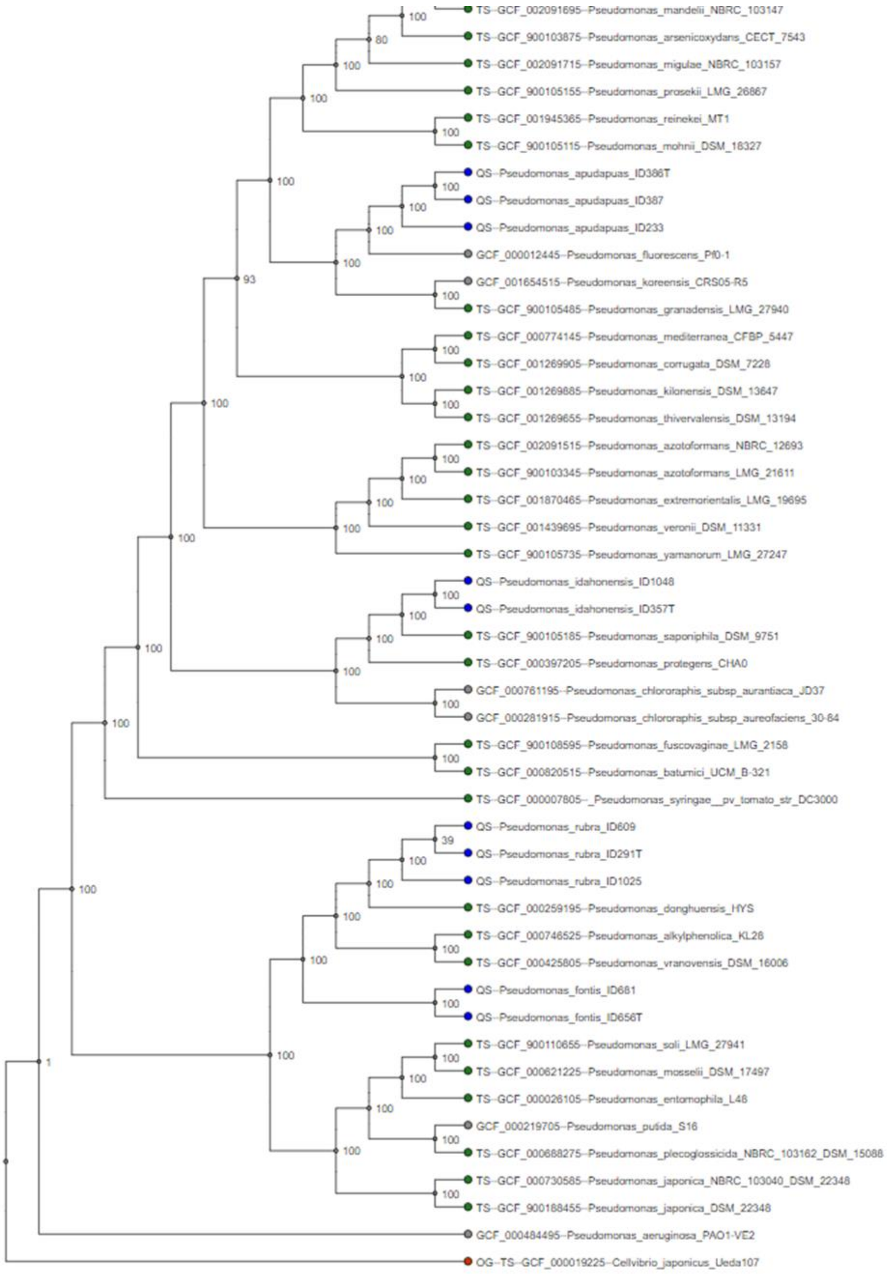

**Supplementary Figure 2: 100 gene MLSA Phylogeny from autoMLST.** Ultra-fast boostrapping was used (100 replicates). Queried genome nodes are in blue, type strain reference genome nodes are in green, non-type strain reference genome nodes are in grey, and the outgroup node is in red. QS – query sequence, TS – type sequence, OG – outgroup.

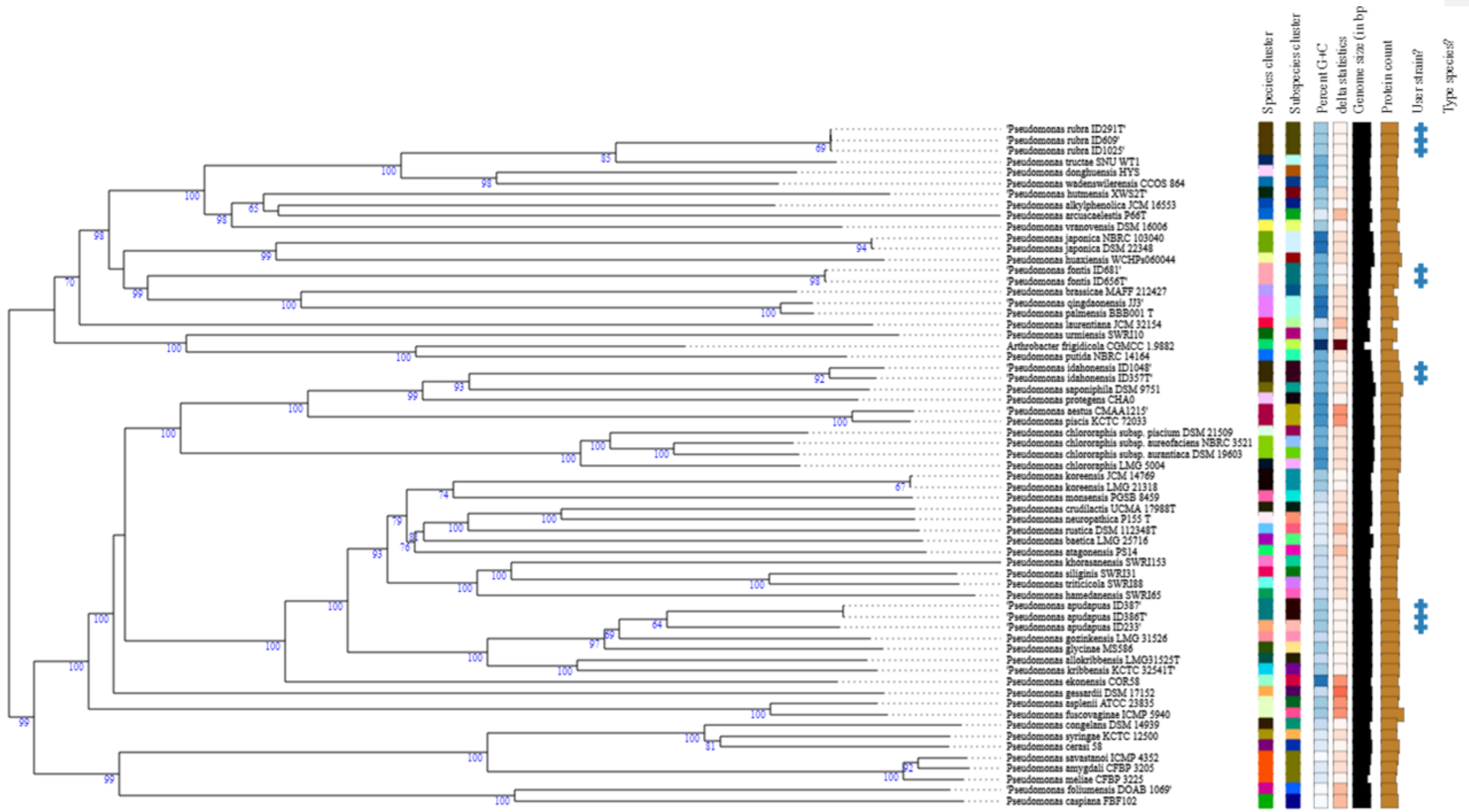


**Supplementary Figure 3: GBDP Phylogram Based on Genomes.** 63 strains were included in this analysis and average bootstrap support was 88.9%. Delta statistic was 0.145.

**Commented [TT1]:** Low quality as PDF - will likely replace with TIFs for submission but trying to avoid making document file size too large

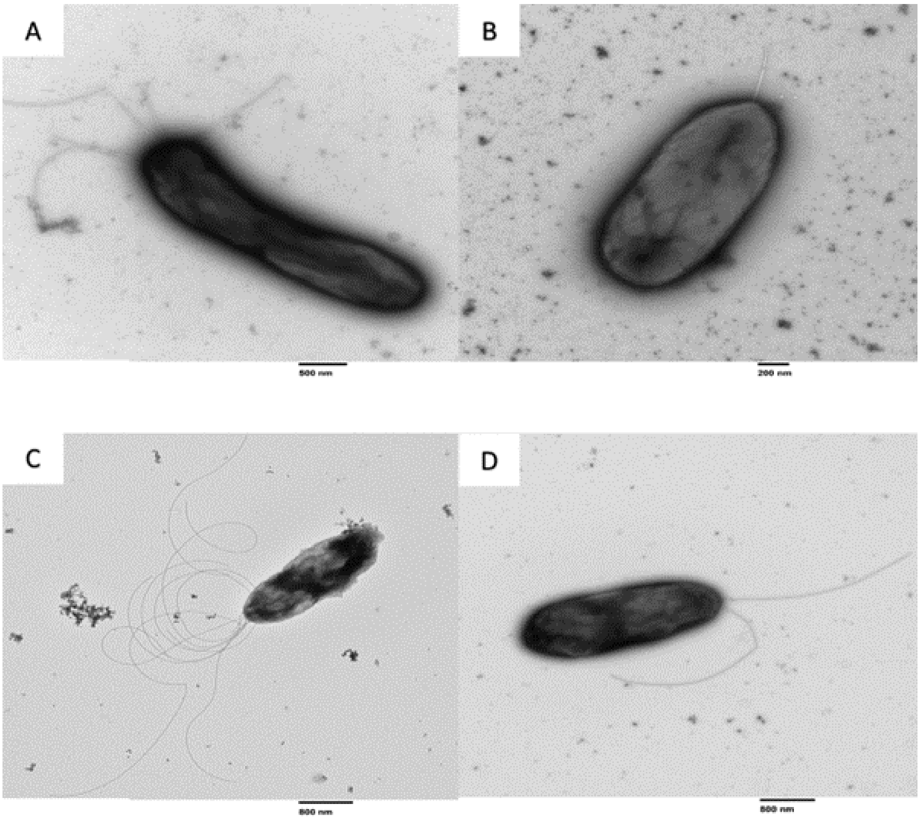


**Supplementary Figure 4: TEM Images of Type Strains.** (A) ID291$^{T}$, (B) ID357$^{T}$, (C) ID386$^{T}$, and (D) ID656$^{T}$. Note that scale bars differs between images

**Supplementary Table 1: Isolation Locations and Conditions.**

| Isolate | Isolation Location | Isolation Medium |
|---|---|---|
| ID233 | Hatchery Raceway Biofilm | TSA |
| ID386[T] | Hatchery Raceway Biofilm | 1:100 TSA |
| ID387 | Hatchery Raceway Biofilm | 1:100 TSA |
| ID291[T] | Natural Rock Biofilm | Nutrient Agar |
| ID609 | Natural Rock Biofilm | R2A |
| ID1025 | Hatchery Raceway Biofilm | Aeromonas Agar |
| ID656[T] | Hatchery Raceway Biofilm | R2A |
| ID681 | Natural Rock Biofilm | R2A |
| ID357[T] | Hatchery Raceway Biofilm | TSA |
| ID1048 | Hatchery Raceway Biofilm | Aeromonas Agar |

**Supplementary Table 2: Genes Used in 100 Gene MLST**

*Separate Excel File will be submitted*

**Supplementary Table 3: Biolog GEN III Results for All Tested Isolates.**

| | 1 | 2 | 3 | 4 | 5 | 6 | 7 |
|---|---|---|---|---|---|---|---|
| **A01 (Negative Control)** | - | - | - | - | - | - | - |
| **A02 (Dextrin)** | - | - | - | w | - | w | - |
| **A03 (D-Maltose)** | - | - | - | - | - | - | - |
| **A04 (D-Trehalose)** | + | - | - | + | - | - | - |
| **A05 (D-Cellobiose)** | - | - | - | - | - | - | - |
| **A06 (β-Gentiobiose)** | - | - | - | - | - | - | - |
| **A07 (Sucrose)** | + | - | - | + | - | - | - |
| **A08 (Turanose)** | - | - | - | - | - | - | - |
| **A09 (Stachyose)** | - | - | - | - | - | - | - |
| **A10 (Positive Control)** | + | + | + | + | + | + | + |
| **A11 (pH 6)** | + | + | + | + | + | + | + |
| **A12 (pH 5)** | + | + | + | + | + | + | + |
| **B01 (D-Raffinose)** | - | - | - | - | - | - | - |
| **B02 (α-D-Lactose)** | - | - | - | - | - | - | - |
| **B03 (D-Melibiose)** | - | - | - | - | - | - | - |
| **B04 (β-Methyl-D-Glucoside)** | - | - | - | - | - | - | - |
| **B05 (D-Salicin)** | - | - | - | - | - | - | - |
| **B06 (N-Acetyl-D-Glucosamine)** | w | - | - | - | - | - | - |

| | | | | | | | |
|---|---|---|---|---|---|---|---|
| **B07 (N-Acetyl-β-D-Mannosamine)** | - | - | - | - | - | - | - |
| **B08 (N-Acetyl-D-Galactosamine)** | - | - | - | - | - | - | - |
| **B09 (N-Acetyl-Neuraminic Acid)** | - | - | - | - | - | - | - |
| **B10 (1% NaCl)** | + | + | + | + | + | + | + |
| **B11 (4% NaCl)** | - | + | - | + | + | - | w |
| **B12 (8% NaCl)** | - | - | - | - | - | - | - |
| **C01 (D-Glucose)** | + | + | + | + | + | + | + |
| **C02 (D-Mannose)** | + | w | - | + | + | - | - |
| **C03 (D-Fructose)** | w | - | - | w | - | - | - |
| **C04 (D-Galactose)** | - | + | - | - | + | - | - |
| **C05 (3-O-Methyl-D-Glucose)** | - | - | - | - | - | - | - |
| **C06 (D-Fucose)** | w | + | - | w | + | - | w |
| **C07 (L-Fucose)** | w | w | w | + | w | - | w |
| **C08 (L-Rhamnose)** | - | - | - | - | - | - | - |
| **C09 (Inosine)** | - | - | - | w | w | - | w |
| **C10 (1% Sodium Lactate)** | + | + | + | + | + | + | + |
| **C11 (Fusidic Acid)** | + | + | + | + | + | + | + |
| **C12 (D-Serine #2)** | + | - | + | w | - | + | + |
| **D01 (D-Sorbitol)** | - | - | - | - | - | - | - |
| **D02 (D-Mannitol)** | - | w | - | w | - | - | - |
| **D03 (D-Arabitol)** | - | - | - | - | - | - | - |
| **D04 (myo-Inositol)** | w | - | - | w | - | - | - |
| **D05 (Glycerol)** | w | - | - | w | - | - | + |
| **D06 (D-Glucose-6-Phosphate)** | - | - | - | - | - | - | - |
| **D07 (D-Fructose-6-Phosphate)** | w | - | - | + | - | - | - |
| **D08 (D-Aspartic Acid)** | - | - | - | - | - | - | - |
| **D09 (D-Serine #1)** | - | - | - | - | - | - | + |
| **D10 (Troleandomycin)** | + | + | + | + | + | + | + |
| **D11 (Rifamycin SV)** | + | + | + | + | + | + | + |
| **D12 (Minocycline)** | + | - | + | + | - | + | - |
| **E01 (Gelatin)** | - | - | - | - | - | - | - |
| **E02 (Gly-Pro)** | - | - | - | - | - | - | - |
| **E03 (L-Alanine)** | - | w | - | - | w | - | + |
| **E04 (L-Arginine)** | w | + | - | + | + | - | + |
| **E05 (L-Aspartic Acid)** | - | - | - | w | - | - | + |
| **E06 (L-Glutamic Acid)** | - | w | + | + | - | + | + |
| **E07 (L-Histidine)** | w | - | + | + | - | + | + |
| **E08 (L-Pyroglutamic Acid)** | - | w | w | w | w | - | - |
| **E09 (L-Serine)** | - | + | w | - | + | w | + |
| **E10 (Lincomycin)** | + | + | + | + | + | + | + |

| | | | | | | | |
|---|---|---|---|---|---|---|---|
| **E11 (Guanidine Hydrochloride)** | + | + | + | + | + | + | + |
| **E12 (Niaproof)** | + | + | + | + | + | + | + |
| **F01 (Pectin)** | w | - | - | w | - | - | - |
| **F02 (D-Galacturonic Acid)** | - | - | - | - | - | - | - |
| **F03 (L-Galactonic Acid-γ-Lactone)** | - | w | - | - | w | - | - |
| **F04 (D-Gluconic Acid)** | + | + | w | + | + | w | + |
| **F05 (D-Glucuronic Acid)** | - | - | - | - | - | - | - |
| **F06 (Glucuronamide)** | w | w | - | w | + | - | w |
| **F07 (Mucic Acid)** | - | + | - | - | + | - | + |
| **F08 (Quinic Acid)** | + | + | - | + | + | - | - |
| **F09 (D-Saccharic Acid)** | - | + | - | - | + | - | - |
| **F10 (Vancomycin)** | + | + | + | + | + | + | + |
| **F11 (Tetrazolium Violet)** | + | + | + | + | + | + | + |
| **F12 (Tetrazolium Blue)** | + | + | + | + | + | + | + |
| **G01 (p-Hydroxy-Phenylacetic Acid)** | + | - | + | + | - | + | - |
| **G02 (Methyl Pyruvate)** | - | - | - | - | - | - | + |
| **G03 (D-Lactic Acid Methyl Ester)** | - | - | - | - | - | - | - |
| **G04 (L-Lactic Acid)** | w | + | w | + | + | w | + |
| **G05 (Citric Acid)** | + | + | - | + | + | - | + |
| **G06 (α-Keto-Glutaric Acid)** | + | w | - | + | + | - | + |
| **G07 (D-Malic Acid)** | - | - | - | - | - | - | + |
| **G08 (L-Malic Acid)** | + | + | + | + | + | + | + |
| **G09 (Bromo-Succinic Acid)** | w | w | - | w | w | w | + |
| **G10 (Nalidixic Acid)** | + | + | + | + | + | + | + |
| **G11 (Lithium Chloride)** | + | - | - | - | + | w | - |
| **G12 (Potassium Tellurite)** | - | + | + | + | + | + | + |
| **H01 (Tween 40)** | + | + | w | + | + | w | + |
| **H02 (γ-Amino-n-Butyric Acid)** | + | + | - | + | w | w | + |
| **H03 (α-Hydroxy-Butyric Acid)** | - | - | - | - | - | - | - |
| **H04 (β-Hydroxy-Butyric Acid)** | - | + | - | w | w | - | + |
| **H05 (α-Keto-Butyric Acid)** | - | - | - | - | - | - | w |
| **H06 (Acetoacetic Acid)** | - | - | - | - | - | - | - |
| **H07 (Propionic Acid)** | w | w | - | + | w | - | + |
| **H08 (Acetic Acid)** | + | + | + | + | + | + | + |
| **H09 (Sodium Formate)** | - | - | - | - | - | - | - |
| **H10 (Aztreonam)** | + | + | + | + | + | + | + |
| **H11 (Butyric Acid)** | - | - | - | - | - | - | - |
| **H12 (Sodium Bromate)** | w | w | - | w | w | - | + |

Organisms: 1, ID1048; 2, ID233; 3, ID291$^T$; 4, ID357$^T$; 5, ID386$^T$; 6, ID609; 7, ID656$^T$.
Symbols: -, negative reaction; w, weak reaction; +, positive reaction.

**Supplementary Table 4: FAME and GC-MS Results for the 4 Proposed Type Strains.**

| | ***Pseudomonas apudapuas* ID386[T]** | ***Pseudomonas idahonensis* ID357[T]** | ***Pseudomonas rubra* ID291[T]** | ***Pseudomonas fontis* ID656[T]** |
|---|---|---|---|---|
| $C_{10:0\ 3\text{-}OH}$ | 5.1 | 7.5 | 11 | 9.2 |
| $C_{12:0}$ | 1.5 | 2 | 1.6 | 2.5 |
| $C_{12:0\ 2\text{-}OH}$ | 6.5 | 5.1 | 5.4 | 4.7 |
| $C_{12:1\ 3\text{-}OH}$ ω7c | 0.3 | 3 | 2.4 | 3.1 |
| $C_{12:0\ 3\text{-}OH}$ | 5.1 | 7.2 | 6.1 | 6 |
| $C_{14:0}$ | 0.4 | 0.2 | 0.5 | 0.3 |
| $C_{15:0}$ | TR | TR | TR | TR |
| $C_{14:0\ 3\text{-}OH}$ | ND | 1.2 | 0.6 | 0.5 |
| $C_{16:1}$ ω7c | 39.6 | 21.7 | 22.2 | 20.4 |
| $C_{16:1}$ ω7t | ND | 3.3 | 6.6 | 4.2 |
| $C_{16:0}$ | 28.6 | 22.8 | 23.9 | 22.7 |
| $C_{17:0}$ cyclo ω7c | 1.2 | 4.6 | 1.9 | 6.4 |
| $C_{17:0}$ | ND | 0.1 | ND | ND |
| $C_{16:0\ 3\text{-}OH}$ | ND | 0.2 | ND | ND |
| $C_{18:1}$ ω7c | 11.2 | 19.3 | 17.2 | 19.3 |
| $C_{18:0}$ | 0.3 | 0.7 | 0.3 | 0.5 |
| $C_{18:1}$ ω7c 11-methyl | ND | 0.5 | 0.2 | ND |
| C10,13-epoxy-11-methyl-octadecadienoate | ND | 0.2 | ND | ND |

ND, Not Detected; TR, Trace Amounts
